# Genome-wide transposon screen of a *Pseudomonas syringae mexB* mutant reveals the substrates of efflux transporters

**DOI:** 10.1101/684605

**Authors:** Tyler C. Helmann, Caitlin L. Ongsarte, Jennifer Lam, Adam M. Deutschbauer, Steven E. Lindow

## Abstract

Bacteria express numerous efflux transporters that confer resistance to diverse toxicants present in their environment. Due to a high level of functional redundancy of these transporters, it is difficult to identify those that are of most importance in conferring resistance to specific compounds. The resistance-nodulation-division (RND) protein family is one such example of redundant transporters that are widespread among Gram-negative bacteria. Within this family, the MexAB-OprM protein complex is highly-expressed and conserved among *Pseudomonas* species. We exposed barcoded transposon mutant libraries in isogenic wild-type and Δ*mexB* backgrounds in *P. syringae* B728a to diverse toxic compounds *in vitro* to identify mutants with increased susceptibility to these compounds. Mutants in genes encoding both known and novel redundant transporters, but with partially overlapping substrate specificities were observed in a Δ*mexB* background. Psyr_0228, an uncharacterized member of the Major Facilitator Superfamily of transporters, preferentially contributes to tolerance of acridine orange and acriflavine. Another transporter located in the inner membrane, Psyr_0541, contributes to tolerance to acriflavine and berberine. The presence of multiple redundant, genomically encoded, efflux transporters appears to enable bacterial strains to tolerate a diversity of environmental toxins. This genome-wide screen in a hyper-susceptible mutant strain revealed numerous transporters that would otherwise be dispensable in these conditions. Bacterial strains such as *P. syringae* that likely encounter diverse toxins in their environment such as in association with many different plant species, probably benefit from possessing multiple redundant transporters that enable versatility to tolerate novel toxicants.

## Introduction

Bacteria, like all living organisms, must tolerate a variety of potentially harmful, chemically diverse molecules present in the environment. While many of these compounds can be degraded to prevent their accumulation to harmful levels within cells, a common means of tolerance of toxins is to export them from the cell. Bacterial efflux transporters can function both to achieve stress tolerance as well as to contribute to virulence by secreting toxins or effectors of various kinds (1, 2). A given bacterial species commonly possesses a wide variety of efflux transporters. It is presumed that they differ in the specificity of toxins that they export. Broad-specificity transporters actively remove toxins from the cell, and their substrates can include heavy metals, solvents, dyes, detergents, antibiotics, as well as certain host-derived products (3–6). An example of such promiscuous transporters are multidrug resistant (MDR) efflux transporters that remove a wide range of structurally diverse chemical compounds from the cell interior (7). While genes encoding these exporters can be found on plasmids, both pathogenic and nonpathogenic bacteria have comparable numbers of chromosomally encoded MDR systems (8). For those bacteria that colonize eukaryotic hosts, MDR efflux pumps not only export antibiotics and other toxic compounds found in their environment, but also host-derived antimicrobial compounds (2). Given that genes encoding efflux transporters are highly conserved between diverse bacteria, their physiological role is likely not involved in resistance to clinically relevant antibiotics (9). A high level of redundancy of transporters exists within a given strain, where they may share many of the same substrates (10, 11). These observations prompt the question of why so many transporters are present in a given bacterium, given the broad substrate ranges of these MDR pumps.

MDR efflux transporters are structurally diverse, being found in at least five distinct protein families: the major facilitator superfamily (MFS), the small multidrug resistance (SMR) family, the multidrug and toxic compound extrusion (MATE) family, the ATP-binding cassette (ABC) superfamily, and the resistance-nodulation-division (RND) family (12, 13). Many of these transporter classes contain both substrate-specific transporters as well as less specific MDR-type efflux pumps (8). For example, characterized MFS transporters in *Escherichia coli* range from highly-specific sugar:proton symporters such as LacY or XylE to multidrug efflux transporters like EmrD (14). Our understanding of efflux-mediated resistance to toxins and antibiotics would benefit from both identifying the MDR transporters from within the larger collection of genomic transporters for a given organism, and also identifying the specific substrates for those transporters.

Multiple transporters, in both the same and different protein families, can share the same substrates (2, 10). This redundancy poses a challenge for their study, as these alternative transporters can mask mutations that disrupt the function of a given pump under investigation (10). The large number of potential MDR transporters in a given strain also makes their investigation by analysis of targeted gene deletion strains laborious. An example of one of the few such intensive studies is that of Sulavik *et al*. (15), who tested the susceptibility of *E. coli* strains with null mutations in 7 known and 9 predicted efflux genes against 35 toxicants. This study, and others (16, 17), identified the RND transporter AcrAB-TolC as the major determinant of intrinsic toxicant resistance in *E. coli*.

Homologs of AcrAB-TolC are common in Gram-negative bacteria, and have been shown to contribute to virulence of plant pathogens as diverse as *Erwinia amylovora, Pseudomonas syringae, Ralstonia solanacearum*, and *Xylella fastidiosa* (18–21). Interestingly, inhibition of efflux pumps by known chemical inhibitors can increase the antimicrobial activity of compounds produced by plants (22–24) and might be considered a plant disease resistance trait. *P. syringae* pv. *syringae* is a plant pathogen that is commonly found both in association with plants as well as in the water cycle (25). These are habitats in which it might be expected to encounter a variety of toxic compounds. MexAB-OprM is the best-characterized AcrAB-TolC homolog present in *Pseudomonas* species. While this protein complex is proposed to secrete the iron-chelating molecule pyoverdine (26), it also contributes significantly to antibiotic resistance (12). In *P. syringae* pv. *syringae* strain B728a, genes in this operon are generally expressed at much higher levels than other RND transporters, both in culture as well as in cells on the leaf surface and in the apoplast (27). However, in *P. aeruginosa*, expression of *mexAB-oprM* is inversely correlated with expression of the related RND transporters *mexEF-oprN* and *mexCD-oprJ* (28). This suggests that regulation of the overall repertoire of efflux transporters is tightly regulated, and that alterations in one or more can lead to compensatory changes in the others. To examine the role of MexAB-OprM homologs and other potential MDR transporters present in strain B728a, we interrogated the fitness of a large library of randomly-barcoded *mariner* transposon mutants (RB-TnSeq) in culture media containing diverse antimicrobial compounds. RB-TnSeq is a modification of TnSeq where each transposon insertion is tagged with a unique 20-nucleotide barcode (29). As transposon insertions are only mapped once for a given library, this reduces the effort to use that library for fitness contributions of genes in multiple conditions. Changes in relative barcode abundance over time are used as a proxy for the relative fitness contribution of a given gene in a given condition. This method can be used to associate bacterial genes with their importance to fitness in different growth conditions, and has been used to improve genome annotations for diverse bacteria (30). Of particular importance for this current study is the ease and scale with which this method could be used to associate genes encoding transporter proteins with their likely substrates. Here, we used RB-TnSeq to identify likely substrates for B728a efflux transporters, with a particular focus on complementary RND homologs of MexAB-OprM.

## Results

### Creation of a barcoded transposon library in B728a Δ*mexB*

To test the role of redundant RND efflux proteins *in vitro*, we created a barcoded *mariner* transposon library in a Δ*mexB* strain. Comparisons of gene fitness in the mutant library with that of the barcoded transposon library in wild-type (WT) strain B728a (31), allowed us to directly test the fitness contributions of B728a transporters in both genetic backgrounds, enabling the complementarity of other transporters with MexAB-OprM to be quantified. The Δ*mexB* transposon library contains 237,285 unique insertion strains, and is a similar size and insertion density to the WT library (Table S1). We hypothesized that insertional mutants in efflux genes in the Δ*mexB* genotype would be less fit than mutants in the WT genotype, particularly when exposed to toxic substrates of MexAB-OprM. The B728a genome encodes 668 predicted transport proteins (32); here we primarily focus on the RND transporters but also investigate members of the other protein families that typically encode MDR efflux transporters.

The *mexB* deletion strain completely eliminates the activity of MexAB-OprM, including inactivating OprM. To determine if the deletion of *mexB* resulted in a polar mutation, we expressed *mexAB* and *mexAB-oprM* under the native promoter in the Δ*mexB* mutant strain. While the Δ*mexB* strain containing the plasmid to express the entire operon was able to tolerate acriflavine and berberine at WT levels, the Δ*mexB* strain expressing only *mexAB* was not (Fig. S1). This indicates that the Δ*mexB* transposon library does not produce a functional OprM.

### Testing the fitness of mutants in the WT and Δ*mexB* transposon libraries *in vitro*

Both mutant libraries were grown in the rich culture medium King’s B (KB) with different antimicrobial compounds; for compounds where the minimum inhibitory concentration (MIC) was known for B728a WT and Δ*mexB* (20), we used ¼ the MIC. We successfully assayed fitness for 16 unique antimicrobial compounds (Table S2). For each gene, fitness is calculated as the log_2_ ratio of barcode abundance following growth in a given condition relative to barcode abundance measured initially at time0. As expected, insertions in the majority of genes did not contribute to fitness as measured by relative barcode abundance in the population, and thus the fitness scores for most genes were close to 0. A mutant with a fitness score of −1 is approximately 50% less abundant relative to the typical strain in the library under that experimental condition.

### Identification of MexAB-OprM substrates

For many substrates, the dominant activity of MexAB-OprM seen in other species was expected to mask any requirement for complementary transporters. We therefore hypothesized that the Δ*mexB* deletion library would unmask the contributions of such alternative transporters to tolerance of various toxins. The majority of antimicrobial compounds tested here are substrates of the MexAB-OprM transporter, and have known MICs (20). Other previously non-investigated compounds include capsaicin, flavone, rifampicin, and rotenone.

For the majority of the antimicrobial compounds tested, disruption of any gene in the MexAB-OprM operon resulted in decreased fitness relative to insertions in the average gene in the WT strain (Fig. 1). No fitness change for insertional strains in this operon was observed when cells were exposed to capsaicin, rifampicin, or rotenone. There was also no fitness change when exposed to ampicillin, a previously-reported substrate (20). For erythromycin, only transposon insertions in *oprM* decreased mutant fitness. Flavone is a likely a substrate of this complex due to the large decreases in fitness comparable to that seen upon exposure to known substrates that accompanied disruptions across the *mexAB-oprM* operon. Using this method, we would expect insertional mutations in *mexA* and *oprM* to be neutral in the Δ*mexB* genetic background. All strains in the Δ*mexB* transposon library indeed were unable to assemble a complete MexAB-OprM complex, since insertions elsewhere in the *mexAB-oprM* operon did not alter fitness. For all antimicrobial compounds tested, *mexA* and *oprM* are dispensable in the Δ*mexB* library, with fitness scores close to 0 (Fig. 1).

**Figure 1.**
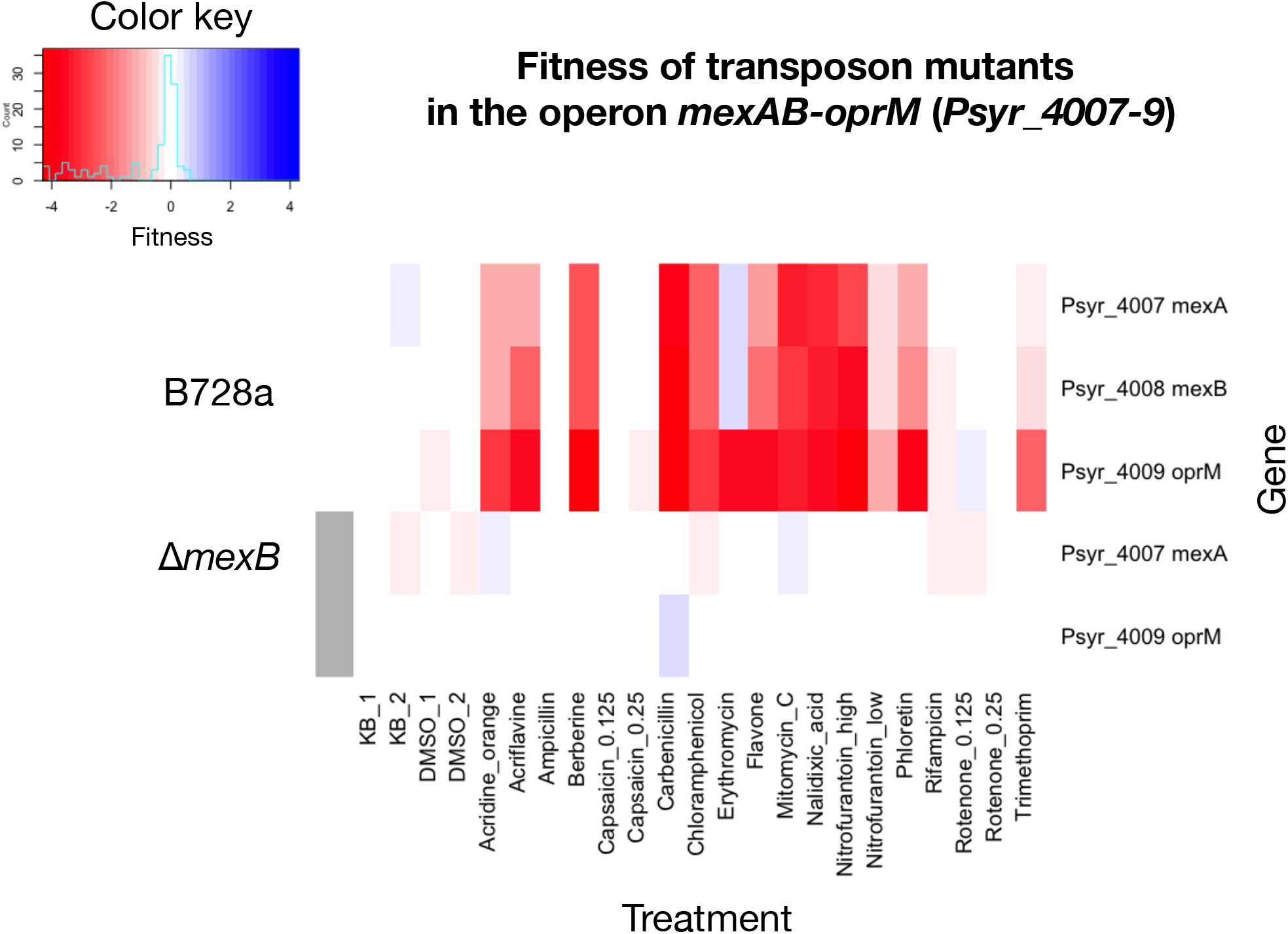
Fitness of transposon insertional mutants in the operon *mexAB-oprM* (*Psyr_4007-9*) in both a WT background and in cells in which *mexAB-oprM* has been disrupted. KB and DMSO are the media and media + solvent control respectively. Fitness is calculated as log_2_ change in relative insertion strain barcode abundance for a given gene.

### Transporters homologous to MexAB contribute to resistance to diverse toxicants

To identify efflux transporters that had the largest contribution to tolerance of various toxicants, we calculated the number of compounds in which genes encoding transporters or transporter components had a fitness score less than −1. RND transporters generally had a larger contribution to toxin tolerance in the Δ*mexB* library than in the WT, but some were important in both backgrounds (Table 1). B728a contains 16 RND transporters, including MexAB-OprM (32). Due to our experimental design that focused on known MexAB-OprM substrates, *mexA* and *mexB* were required for competitive fitness in the presence of 10 compounds. The outer membrane transporter OprM is likely shared with additional efflux pumps, and so is required for tolerance of an additional 3 compounds.

**Table 1.**
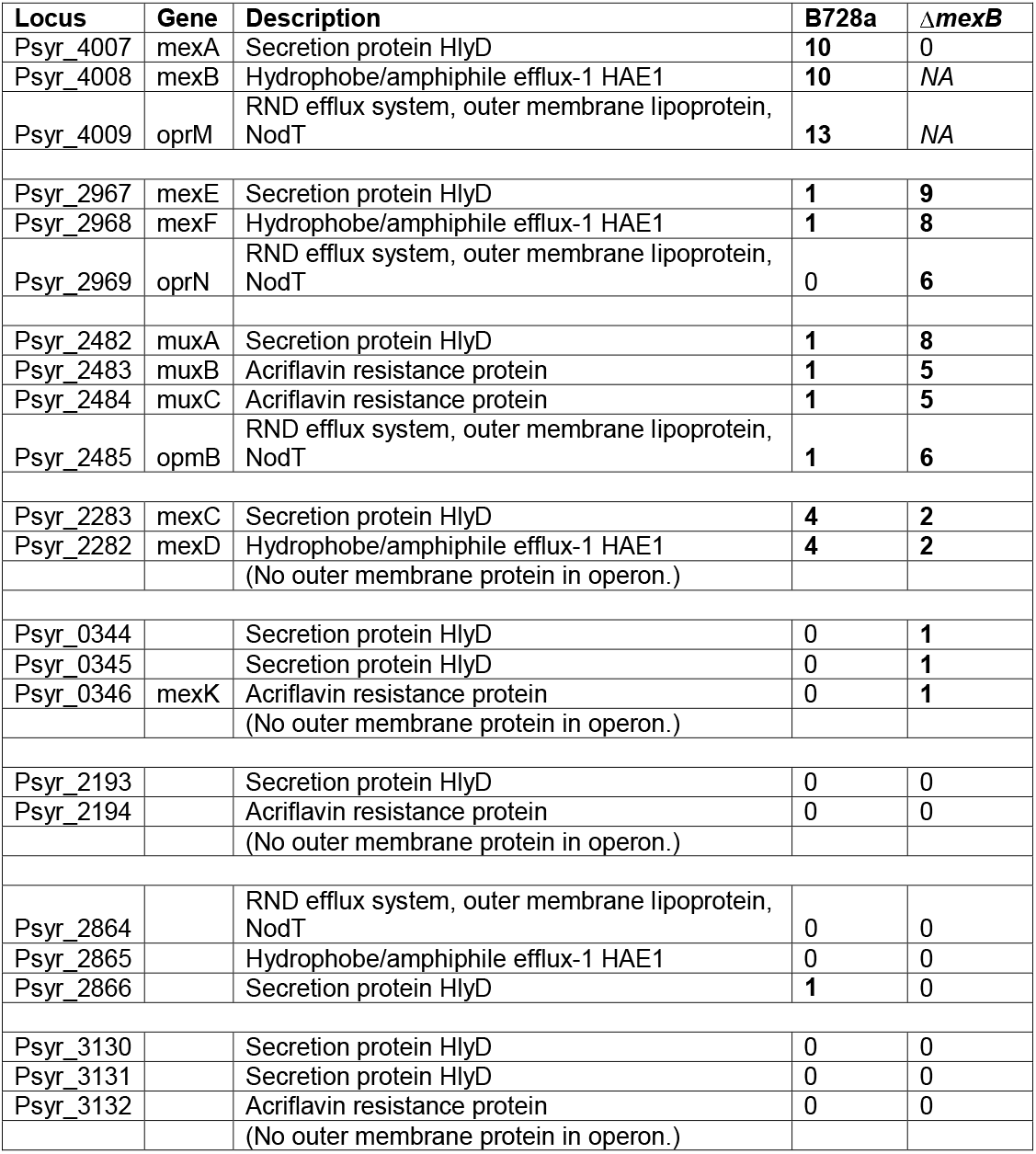
RND operons homologous to *mexAB-oprM* that likely contribute to multidrug resistance. The number of experimental conditions where they have a significant contribution to fitness (fitness value less than −1) is shown for mutants in a WT and Δ*mexB* background. A total of 23 treatments were examined, including 2 KB controls, 2 DMSO controls, and 16 unique compounds (3 at two concentrations).

In addition to MexAB, four homologous RND transporters contributed substantially to tolerance of several toxicants (Table 1). The *mexEF-oprN* operon however, was only required for tolerance of these compounds in the absence of MexB (Fig. 2). MexEF-OprN is likely redundant with MexAB-OprM for the substrates acridine orange, acriflavine, berberine, chloramphenicol, flavone, nalidixic acid, nitrofurantoin, and phloretin. The lack of a phenotype for mutants in the WT background is consistent with the likely subsidiary role of this transporter for these shared substrates, and therefore masking by the more highly expressed MexAB-OprM under these conditions.

**Figure 2.**
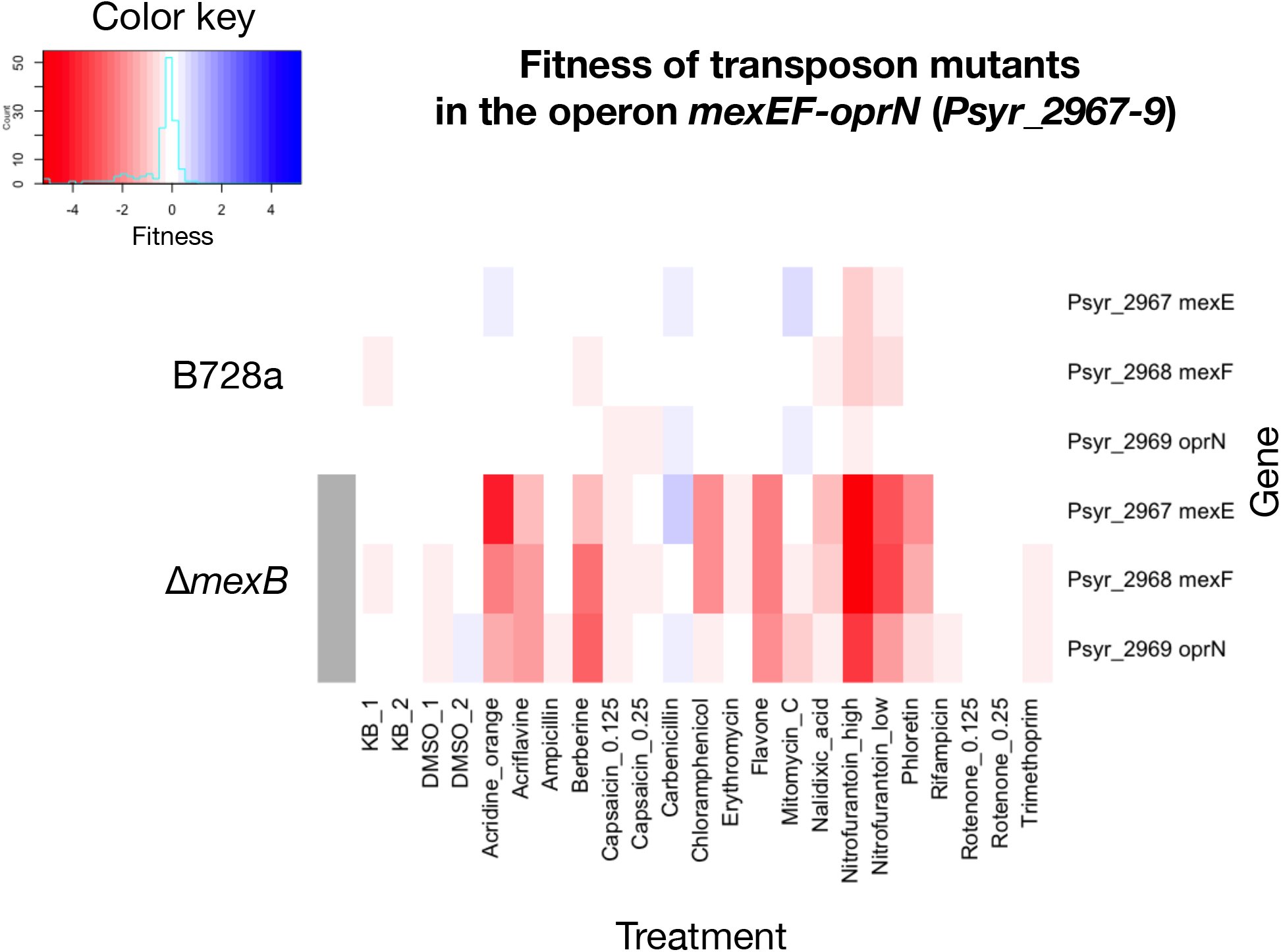
Fitness of transposon insertional mutants in the operon *mexEF-oprN* (*Psyr_2967-9*) in both a WT background and in cells in which *mexAB-oprM* has been disrupted. Fitness is calculated as log_2_ change in relative insertion strain barcode abundance for a given gene.

Psyr_2483-5 is likely redundant with MexAB-OprM for the substrates acriflavine, berberine, carbenicillin, chloramphenicol, erythromycin, and phloretin, with decreased fitness for disruptions in the Δ*mexB* genetic background (Fig. 3). Psyr_2483-5 independently contributes to resistance to rifampicin. This apparent operon encodes an unnamed RND transporter, and is most homologous to the MuxABC-OpmB complex in other Pseudomonads, such as *P. aeruginosa* PAO1 (Table S3).

**Figure 3.**
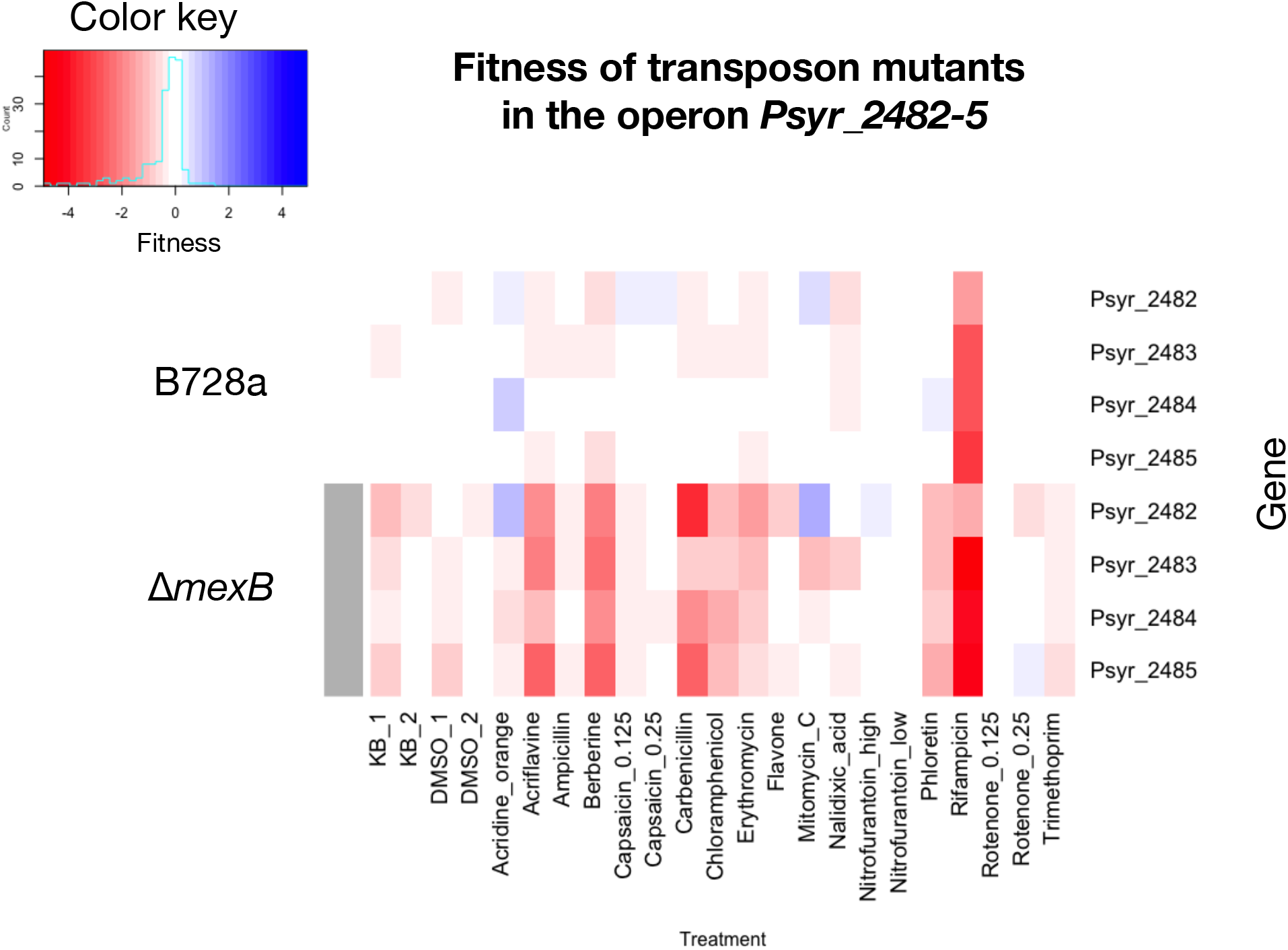
Fitness of transposon insertional mutants in the operon *muxABC-opmB* (*Psyr_2482-5*) in both a WT background and in cells in which *mexAB-oprM* has been disrupted. Fitness is calculated as log_2_ change in relative insertion strain barcode abundance for a given gene.

MexCD (*Psyr_2282-3*) was required for full competitive fitness in the presence of acriflavine, berberine, erythromycin, and nalidixic acid (Fig. S2). Importantly, these phenotypes could be seen in both the WT and Δ*mexB* backgrounds. Negative fitness scores for insertional mutants in this operon were, however, greater in the WT background, although this may have been at least partially due to the higher toxicant concentrations used in testing the mutants in the WT background than that of the Δ*mexB* library. This operon does not encode an outer membrane protein, and a gene for such a required component is likely located elsewhere in the genome.

Of the compounds tested here, Psyr_0344-6 only contributed to tolerance of phloretin in the Δ*mexB* background (Fig. 4). Interestingly, the disruption of any gene in this operon resulted in a mutant strain that was more fit than mutants of other genes when exposed to acriflavine, berberine, or nalidixic acid, but only if *mexB* was also absent. This operon likely requires an unknown outer membrane protein located elsewhere in the genome.

**Figure 4.**
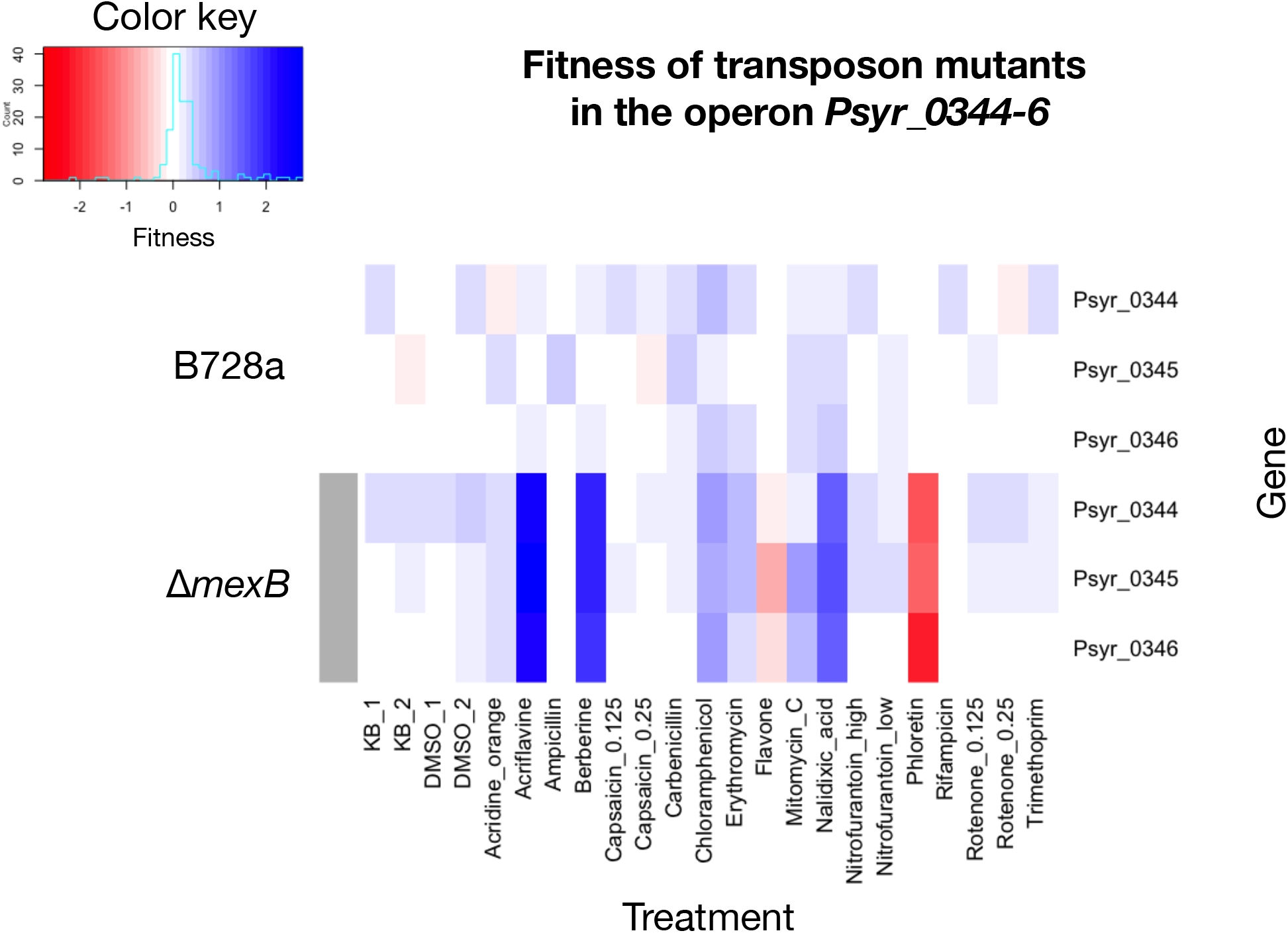
Fitness of transposon insertional mutants in the operon *Psyr_0344-6* in both a WT background and in cells in which *mexAB-oprM* has been disrupted. Fitness is calculated as log_2_ change in relative insertion strain barcode abundance for a given gene.

### Inner membrane transporters appear more substrate specific

The same computational analysis was used to interrogate the MFS and other inner membrane transporters to identify those with fitness contributions in the presence of various toxic compounds in either or both genetic backgrounds. Among 68 MFS transporters examined, only Psyr_0228 contributed to tolerance of any compounds. Psyr_0228 contributed to competitive fitness in both acriflavine and acridine orange (Fig. 5). Interestingly, this gene contributed to acriflavine resistance even in a WT background, indicating that its role was independent of *mexB*. In contrast, this gene contributed to acridine orange resistance only in the Δ*mexB* genotype. Disruption of the gene encoding the SMR transporter Psyr_0541 strongly decreased fitness in berberine and mildly decreased fitness in acriflavine, independent of the *mexB* genotype (Fig. 6). In the Δ*mexB* strain, disruption of *Psyr_0541* resulted in a mild susceptibility to carbenicillin (Fig. 6).

**Figure 5.**
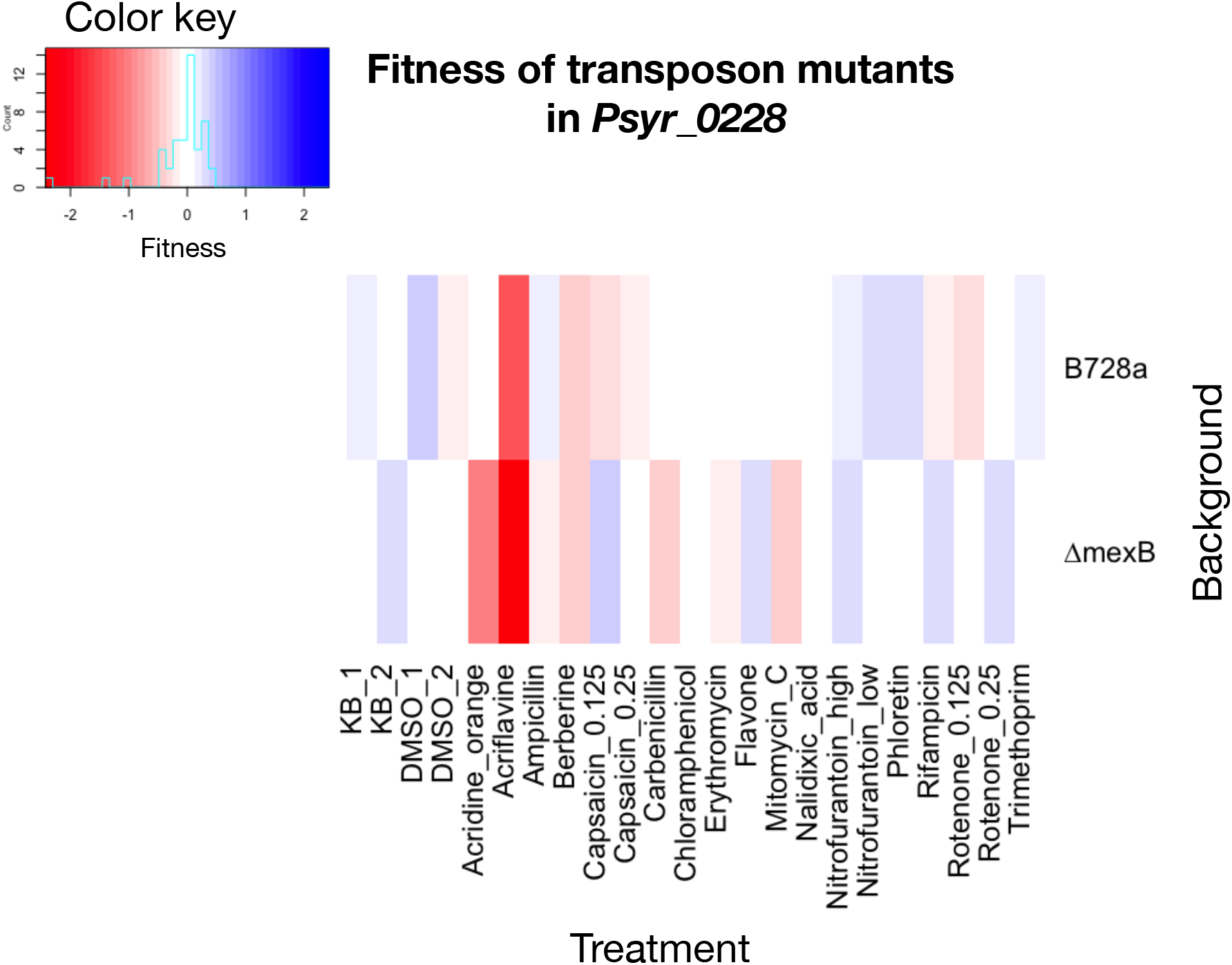
Fitness of transposon insertional mutants in the Major Facilitator Superfamily transporter gene *Psyr_0228* in both a WT background and in cells in which *mexAB-oprM* has been disrupted. Fitness is calculated as the log2 change in relative insertion strain barcode abundance for a given gene.

**Figure 6.**
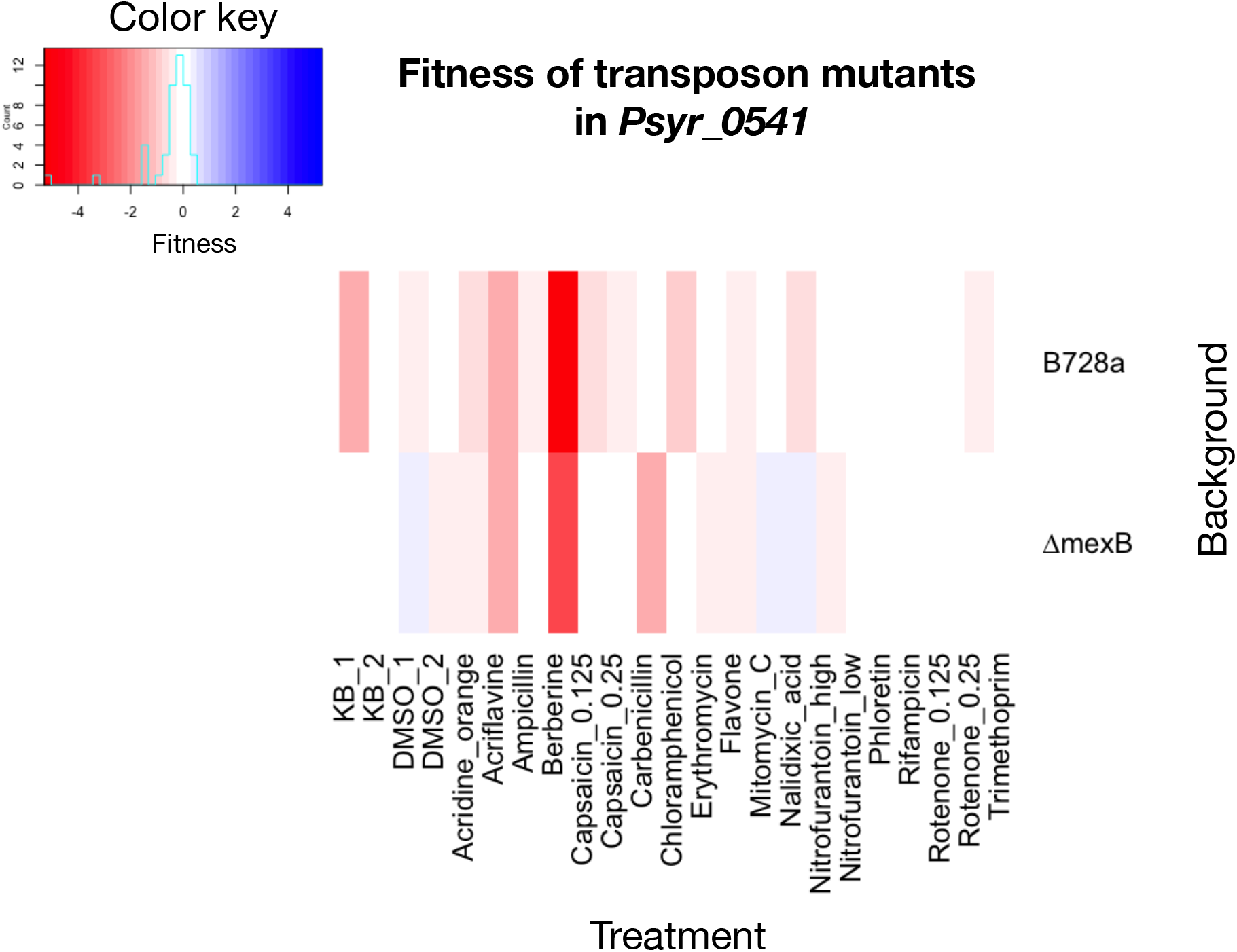
Fitness of transposon insertional mutants in the SMR gene *Psyr_0541* in both a WT background and in cells in which *mexAB-oprM* has been disrupted. Fitness is calculated as the log_2_ change in relative insertion strain barcode abundance for a given gene.

### Several ABC transporters contribute to growth in rich media

Most genes annotated as encoding ABC transporter subunits were putative amino acid or carbohydrate transporters. In the conditions tested, several ABC transporters were required for competitive fitness in the rich media controls. For example, insertions in *Psyr_0917-8*, encoding the polysaccharide permease ABC transporter RfbAB-2 strongly decreased fitness in all conditions, including the controls containing only KB medium, in both the WT and Δ*mexB* genotypes (Fig. S3). RfbAB-2 appears nearly essential for growth in KB. Genes encoding a putative peptide ABC transporter, *Psyr_1754-9*, were required for competitive fitness under all conditions, including the KB controls, but only in a Δ*mexB* mutant background (Fig. S4).

### Overlapping substrate specificities between transporters

The MDR efflux transporters tested here displayed a range of apparent substrates, with varying degrees of overlap with MexAB-OprM (Fig. 7). Compounds such as acriflavine, berberine, and phloretin are probable substrates of multiple RND transporters. MexCD was required for tolerance of fewer compounds tested than MexAB, MexEF, or MuxABC. As hypothesized, the inner membrane transporters Psyr_0228 and Psyr_0541 contributed to resistance to only a few toxicants. However, for the compounds examined here, no other inner membrane transporters contributed to competitive fitness.

**Figure 7.**
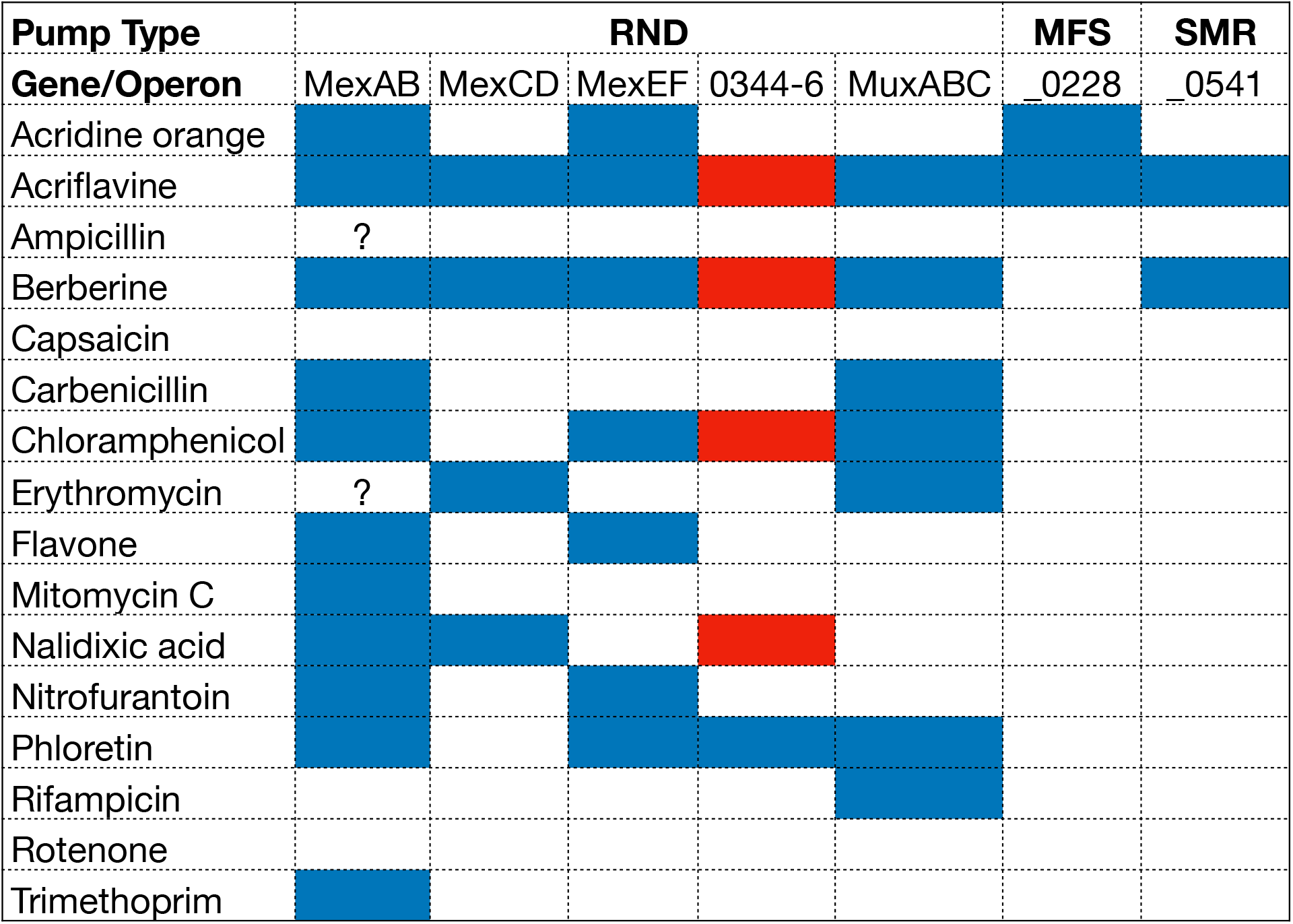
Likely substrates of *P. syringae* B728a multidrug resistance RND transporters, the MFS transporter Psyr_0228, and the SMR pump Psyr_0541. Blue cells indicate the compound is an apparent substrate of that transporter, with insertional mutants of a given gene having decreased fitness (fitness score less than −1) in the WT and/or the Δ*mexB* genetic backgrounds. Red cells indicate an increased fitness of insertional mutants in this operon.

### Zone of inhibition assays to test predicted mutant phenotypes

The fitness contributions of the various transporters were all conducted at a given concentration of a particular toxicant. The lower fitness seen for various mutants suggested a hypersensitivity to that compound. To demonstrate however that disruption of these transporters reduced the concentration of the toxicant at which any growth could occur we constructed targeted deletion mutants of the substrate binding proteins MexF, MuxB, and MexK (Psyr_0346). We used zone of inhibition (ZOI) assays to provide semi-quantitative estimates of the MIC of the toxicants. The Δ*mexB*Δ*mexF* double mutant was significantly more susceptible to berberine, phloretin, and rifampicin than the WT and either single deletion strain (Fig. 8a). The Δ*mexB*Δ*muxB* double mutant was significantly more sensitive to acriflavine and rifampicin than the WT and either single deletion strain (Fig. 8b). As suggested by positive fitness values observed in mixture studies, the Δ*mexBmexK* double mutant exhibited significantly decreased susceptibility to acriflavine and berberine than the Δ*mexB* mutant (Fig. 8c).

**Figure 8.**
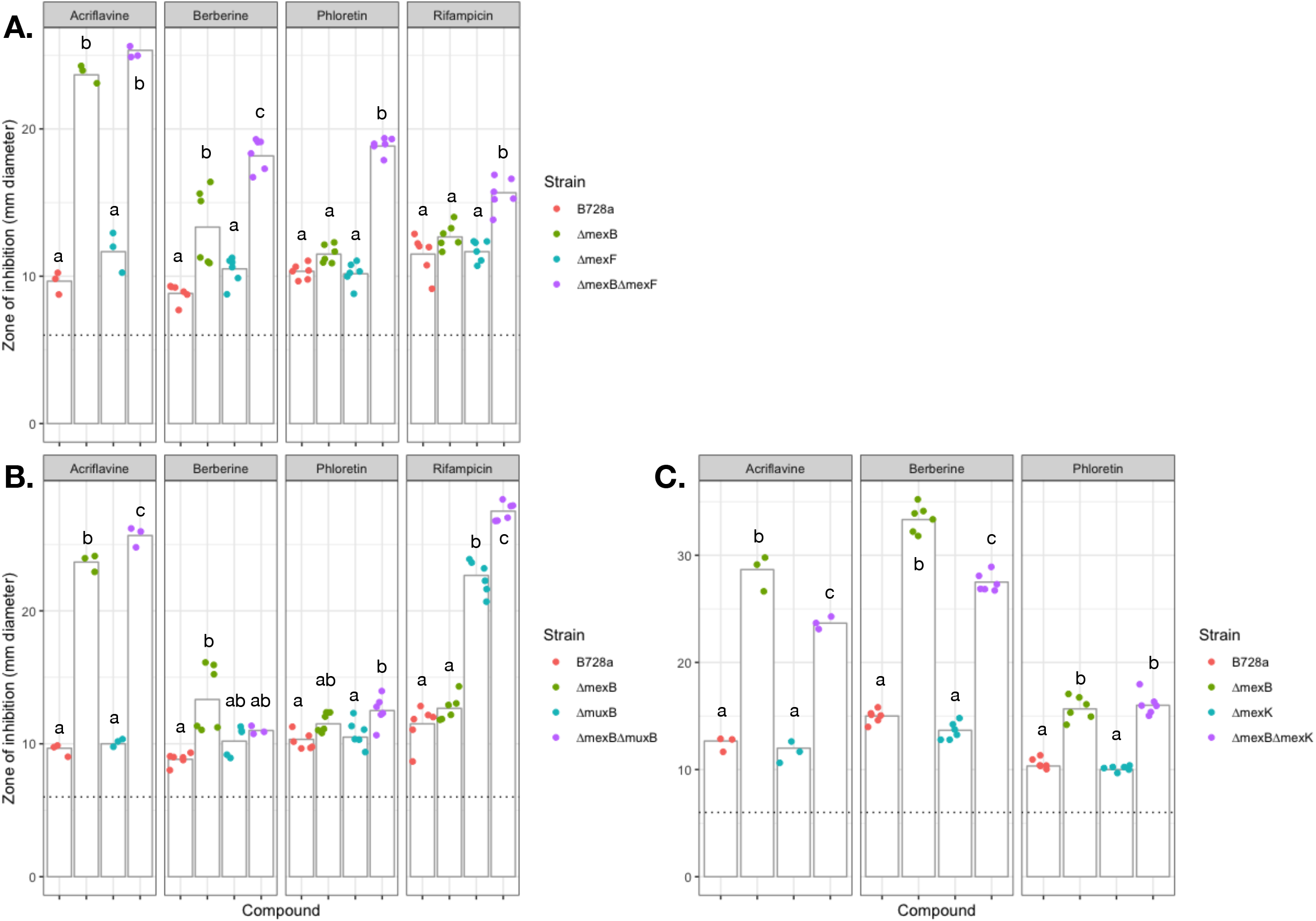
Zone of growth inhibition assays to test antibiotic sensitivity of B728a efflux pump deletion mutants Δ*mexB*Δ*mexF* (A), Δ*mexB*Δ*muxB* (B), and Δ*mexB*Δ*mexK* (C). Mean values are displayed as column height. For each compound tested, means marked with the same letter do not differ at p = 0.01 using Tukey’s HSD test. The lower limit of measurement is 6 mm (dotted line).

## Discussion

Bacteria encounter and must tolerate diverse and potentially toxic molecules in the various habitats they occupy. While efflux transporters are generally thought to play a major role in the tolerance of toxins, it is still unclear why bacterial genomes encode such a large number of MDR transporters. With high expression and wide substrate ranges, RND efflux transporters such as AcrAB-TolC and MexAB-OprM are essential for environmental survival and colonization of eukaryotic hosts (33). However, for these same reasons, these transporters are thought to mask the contributions to resistance of homologous RND transporters (15). Prompting our investigation in *P. syringae*, studies have shown that expression of multiple alternative transporters is increased when these primary RND pumps are disrupted (28, 34). Our studies of these alternative and redundant pumps, has benefited from a method that has previously been successfully employed that relies on the use of hypersusceptible pump mutants, to unmask redundant transporters (35–37).

B728a contains a diverse array of transporters, many with likely roles in multidrug resistance (32). The MexAB-OprM complex has been shown to be important for tolerance of several *P. syringae* strains to a variety of antibiotics and other toxins, as well as virulence during plant colonization (20). However, it has remained a challenge to associate particular transporters with their substrates, especially for host-produced compounds. One strategy to overcome the masking effect of such dominant transporters is either to genetically disrupt genes encoding MDR efflux transporters, as discussed previously, or chemically inhibit them (22, 24, 38). Here, we were able to use a transposon library constructed in a Δ*mexB* strain to reveal the role of several partially redundant RND efflux pumps. While several of these pumps appeared completely masked by the role of MexAB-OprM under the in vitro conditions used here, some were complementary to MexAB-OprM, contributing incrementally to tolerance of various toxicants.

Among those transporters with an incremental role in tolerance were *mexEF-oprN* and *muxABC-opmB*. These transporters however, were only important for a subset of the MexAB-OprM substrates tested. We expected MexEF-OprN to be important in the Δ*mexB* genotype for two main reasons. First, previous transcriptional measurements in *P. syringae* strain B728a show *mexEF-oprN* is generally the second-highest expressed RND transporter after *mexAB-oprM* under a variety of conditions in culture and *in planta* (27). Second, over-expression of *mexEF-oprN* in *P. aeruginosa* strains lacking *mexAB-oprM* increased resistance to several antibiotics (11). MuxABC-OpmB is an unusual RND operon because it is expected to have two inner membrane components. This transporter was only recently characterized in *P. aeruginosa* (39), and is homologous to the *E. coli* complex MdtABCD (also annotated as YegMNOB) (35, 36). Rifampicin was not previously known to be a substrate of any Mex pumps (11), but full resistance appears to require the MuxABC-OpmB complex.

Of the eight RND operons likely involved in multidrug resistance, four do not contain genes encoding outer membrane proteins. Transporters such as these likely require the presence of outer membrane proteins encoded by genes located elsewhere in the genome. The archetypal outer membrane protein TolC and its homologs (such as OprM and OprN in *Pseudomonas* species) can couple to many different transporters, and apparently do not play a role in substrate specificity (40). Since the construction of the unmarked Δ*mexB* strain also disrupted the *oprM* gene, this may have reduced the functionality of transporters MexCD, Psyr_0344-6, Psyr_2193-4, and Psyr_3130-2 in that strain if they require OprM for maximum functionality. Combined with the minor roles of these other transporters relative to that of MexAB-OprM, this would potentially explain the lack of fitness cost upon their disruption in both the WT and Δ*mexB* genotypes. We should emphasize however that nearly all of the compounds tested here were known MexAB-OprM substrates. In such a scenario, it might be expected that MexAB-OprM would play a dominant role in tolerance to such compounds. These other transporters, however, probably are not masked by the role of MexAB-OprM for many other compounds and conditions in which the cells might encounter them. Very few studies have addressed the breath of the substrate range for MexAB-OprM beyond that of clinically important antibiotics, and given the wide variety of other toxic natural products that bacteria such as *P. syringae* would encounter in its myriad of habitats, it seems likely that many would be found for which these other efflux transporters might play a dominant role. Indeed, evidence for such a role for Psyr_0228 in the tolerance of acriflavin was seen here. It should prove fruitful to test this hypothesis further by the testing of these transposon libraries for their tolerance of diverse additional compounds. Not only will this provide further evidence of the constraints of MexAB-OprM in the chemical ecology of *P. syringae*, but also provide contextual evidence for the retention of the myriad of other efflux pumps in this species that presumably remain under selection, suggesting the importance in at least some settings encountered by this cosmopolitan bacterium. While we observed redundancy in efflux transporter function, seen in the overlapping substrate range of several RND transporters, it may simply benefit the cell to have multiple mechanisms of resistance for a given toxicant.

In Gram-negative bacteria, inner membrane transporters such as those in the MFS can function cooperatively with outer membrane transporters (like RND transporters) (10). This has been proposed to synergistically increase overall toxin resistance for the cell (11). Together with the high level of substrate promiscuity for most MDR RND transporters, we reasoned that inner membrane transporters might contribute to substrate specificity. This would prevent unnecessary efflux, attenuating the proton motive force. If this were true, we would expect to observe that inner membrane transporters would each contribute strongly to resistance of only a few toxicants. For Psyr_0228, this appeared to be the case, which contributed only to the tolerance of acridine orange and acriflavine, two molecules that are structurally similar (Fig. S5). (Technically three molecules, as acriflavine is typically sold as a mixture of the related acriflavine + proflavine). The SMR protein Psyr_0541 is homologous to the quaternary ammonium compound-resistance protein QacE in *P. fluorescens* SBW25 and QacH in *P. aeruginosa* PA14 (41). Both acriflavine and berberine are quaternary ammonium compounds, and it is noteworthy that *Psyr_0541* was found to be necessary for tolerance to both of these compounds. Insertional mutations at this locus were particularly susceptible to berberine. The inner membrane transporters Psyr_0228 and Psyr_0541 appear to share some substrates with MexAB-OprM. More information is required to determine to what extent these transporters function cooperatively if at all. In addition, it is possible that these transporters interact with molecules that are not substrates of MexAB-OprM, potentially with the assistance of outer membrane transporters other than OprM.

Efflux transporters are typically easily detected and annotated computationally due to their sequence homology and transmembrane domains. Determining their substrates is another matter. Here we show that RB-TnSeq can be used to characterize the substrates of MDR efflux transporters, using a range of structurally diverse antimicrobial compounds. This method is sensitive because it measures competitive fitness, rather than simple inhibition of growth, a phenotype important to bacteria in complex environments where they face competition with other microbes. RB-TnSeq is also very cost effective once transposon mutants are initially mapped, since amplification of the barcodes in the same library enables up to 96 experiments run on a single Illumina sequencing lane (29). This high throughput makes it practical to readily test a large number of conditions and compounds. Characterizing MDR transporter redundancy is essential to our understanding of not just clinical antibiotic resistance, but also the role of these abundant proteins for bacterial survival in diverse environments.

Using RB-TnSeq in a hyper-susceptible mutant strain allowed us to examine the role of alternative transporters that might not otherwise be active or discernible. Furthermore, plant-produced compounds are often not as toxic to bacteria as common antibiotics produced by bacteria or fungi, and this has been hypothesized to be due to the activity of efflux transporters (22). There is some evidence that plants can produce MDR pump inhibitors in addition to antimicrobial compounds (42), and this reinforces the importance of multi-“drug” resistance transporters in tolerating diverse plant-produced and other naturally occurring antimicrobial molecules. Multiple-component, outer membrane transporters are apparently highly redundant, while inner membrane transporters can be more substrate specific. This may help explain the abundance of partially redundant MDR efflux transporters found in many bacteria, especially those strains typically found in environments with exposure to diverse toxic chemicals.

## Materials and Methods

### Bacterial strains and growth media

*P. syringae* pv. *syringae* B728a was originally isolated from a bean leaf (*Phaseolus vulgaris*) in Wisconsin (43). The complete genome sequence for B728a is available on NCBI GenBank as accession CP000075.1 (44). B728a and derivative mutant strains were grown on King’s B (KB) agar or in KB broth (45), at 28°C. *E. coli* strains S17-1, TOP10, and XL1-Blue were grown on LB agar or in LB broth at 37°C. When appropriate, the following antibiotics were used at the indicated concentrations: 100 μg/ml rifampicin, 50 μg/ml kanamycin, 15 μg/ml tetracycline, 100 μg/ml spectinomycin, and 40 μg/ml nitrofurantoin.

### Complementation of the *mexB* deletion strain

Expression constructs of *mexAB* and *mexAB-OprM* were constructed through PCR amplification of this partial or complete operon, as well as 1161 bp of upstream sequence that would include potential promoter regions. These PCR products were ligated into the XbaI and EcoRI restriction enzyme sites of the plasmid p519*ngfp* (46). The ligation mixture was subsequently transformed into chemically competent *E. coli* XL1-Blue. Plasmids were confirmed to contain the correct insertion sequences by Sanger sequencing, and then electroporated into the *E. coli* donor strain S17-1. Plasmids were introduced into B728a WT and B728a Δ*mexB* by conjugation on KB overnight, and then selected for 3 days on KB containing kanamycin and nitrofurantoin (*E. coli* counter selection).

### Construction of bar-coded transposon libraries

Construction of the B728a WT barcoded transposon insertion library was described in (31). The Δ*mexB* library was constructed in a similar fashion. For the Δ*mexB* library, we first removed the kanamycin resistance cassette from the B728a Δ*mexB* deletion strain. The pFLP2Ω plasmid containing the Flp recombinase and a spectinomycin resistance cassette (47) was introduced into the Δ*mexB* strain by conjugation. Exconjugants were selected on spectinomycin. Colonies were screened for the loss of kanamycin resistance by plating, and the loss of the kanamycin cassette in sensitive colonies was confirmed by PCR. The pFLP2Ω plasmid was cured from the deletion strain by several overnight passages in KB containing rifampicin only, and spectinomycin sensitivity was confirmed by plating. The rifampicin resistant Δ*mexB* deletion strain was used as the recipient for conjugation with the barcoded *mariner* transposon library using the same protocol.

### *In vitro* growth of the library

Aliquots of the transposon libraries that were stored at −80°C were removed from cold storage, thawed, and inoculated into 25 ml fresh KB with 100 μg/ml kanamycin and grown for approximately 7 hours at 28°C with shaking until the culture reached mid-log phase, OD_600_ 0.5 - 0.7. For time0 samples, 1 ml aliquots were pelleted by centrifugation and the pellets were frozen at −20°C until DNA purification. All *in vitro* experiments were performed in 24-well plates, containing 1 ml total volume per well. Each compound used was added to KB containing 100 μg/ml kanamycin to a final volume of 950 μl. Cell suspensions (50 μl) were added to each well last. Libraries were grown overnight (15 hours) at 28°C with shaking. The cells from each well were then pelleted and frozen until DNA purification.

### DNA isolation and library preparation

DNA from frozen pellets was isolated using the Qiagen DNeasy Blood & Tissue Kit according to manufacturer’s instructions. Cell lysis was performed at 50°C for 10 minutes as per manufacturer instructions. Purified genomic DNA was measured on a nanodrop device and 200 ng of total DNA was used as a template for DNA barcode amplification and adapter ligation as established previously (29). For each time0 sample, two separately purified DNA samples were sequenced as technical replicates.

### Sequencing and fitness data generation

Barcode sequencing, mapping, and analysis to calculate the relative abundance of barcodes was done using the RB-TnSeq methodology and computation pipeline developed by Wetmore *et al*. (29); code available at bitbucket.org/berkeleylab/feba/. Briefly, TnSeq was used to map the insertion sites and associate the DNA barcodes to these insertions. Fitness values for each gene are calculated as the log_2_ ratio of relative barcode abundance following library growth in a given condition divided by the relative abundance in the time0 sample. Barcode counts were summed between replicate time0 samples. For analysis, genes must have adequate coverage in the time0 sample: at least 3 reads per strain and 30 reads per gene (29). Fitness values are normalized across the genome so the median gene has a fitness value of 0. All experiments described herein passed previously described quality control metrics (29). Experimental fitness values are publically available at fit.genomics.lbl.gov.

### Identification of efflux pumps and assessment of fitness

We focused our initial analysis on genes that are homologous to *mexAB-oprM*. In the Transport Database (membranetransport.org) (32), B728a is annotated as containing 16 RND transporters. We manually curated this list of genes to focus on those operons potentially involved in “multidrug resistance”: namely known *mex* genes or operons containing an “acriflavin resistance” or “hydrophobe/amphiphile efflux-1” and HlyD secretion protein. We filtered out 2 pseudogenes and 6 genes with known or likely unrelated function: protein export proteins SecD and SecF (Psyr_1230-1), syringomycin efflux (Psyr_2622), and heavy metal efflux pump CzcA (Psyr_4803). *Psyr_1128* is adjacent to a heavy metal (Cu/Ag) two-component system. *Psyr_1701* is a hypothetical protein in the syringolin A biosynthesis gene cluster. For the remaining 8 RND operons, we compared fitness values for all genes within each operon across all experiments that passed quality control in both libraries. We focused on pumps where multiple encoding genes in the same operon contributed strongly to competitive fitness (fitness score < −1). As a negative control, the 8 genes with expected functions unrelated to drug resistance were confirmed to not be required in any conditions tested (fitness > −0.5 in both libraries). We performed the same analysis using 68 predicted MFS genes (a mixture of potential drug resistance transporters as well as diverse sugar transporters), 210 ABC transporter genes, the SMR gene *Psyr_0541*, and *norA* (*Psyr_0073*). Heatmaps were plotted in R (48) using the gplots package, version 3.0.1.1 (49).

### Construction of targeted deletion mutants

Deletion strains were constructed using an overlap extension PCR protocol as described previously (50). Briefly, 1 kb DNA fragments upstream and downstream of the genes of interest were amplified along with a kanamycin resistance cassette from pKD13 (51). These three fragments were joined by overlap extension PCR. The resulting fragment was blunt-end ligated into the SmaI site of pT*sacB* (52), and transformed into the *E. coli* subcloning strains TOP10 or XL1-Blue, and then the *E. coli* conjugation donor strain S17-1. This suicide plasmid was conjugated into B728 on KB overnight, and then selected for 3 days on KB containing kanamycin and nitrofurantoin. Putative double-crossover colonies that were kanamycin resistant and tetracycline sensitive were selected for screening using external primers and further confirmed by PCR and Sanger sequencing.

### Zone of inhibition assays

Zone of inhibition assays were used to test deletion mutants for altered susceptibility to selected toxicants *in vitro*. Strains were grown on KB agar containing rifampicin overnight, resuspended in 10 mM KPO_4_, and diluted to an OD_600_ of 0.1. 200 μl of the cell suspension was spread on each plate. 6 mm diameter paper discs were used to absorb antibiotics at the following concentrations: acriflavin, 20 μl of 3 mg/ml H_2_O; berberine, 30 μl of 50 mg/ml DMSO; flavone, 15μl of 20 mg/ml DMSO; and phloretin, 30 μl of 30 mg/ml DMSO. 3 replicate discs were used per plate. Plates were incubated at 28°C overnight, and the diameter of the clearance zone was measured. Graphs were plotted in R using the ggplot2 package version 3.1.1 (53).

## Acknowledgements

We thank Morgan Price for assistance with RB-TnSeq sequence analysis, and Jayashree Ray for assistance mapping the insertions sites of the B728a WT and Δ*mexB* transposon libraries. We thank Olga Smelik for experimental assistance. Capsaicin, flavone, and rotenone were kindly provided by Cat Adams and Dr. Thomas Bruns at UC Berkeley. Funding for Tyler Helmann was partially provided by the Arnon Graduate Fellowship and the William Carroll Smith Fellowship. This work used the Vincent J. Coates Genomics Sequencing Laboratory at UC Berkeley, supported by NIH S10 OD018174 Instrumentation Grant.

## Notes

http://fit.genomics.lbl.gov

