## Supplemental Data for "Genome-wide transposon screen of a *Pseudomonas syringae mexB* mutant reveals the substrates of efflux transporters"

**Supplementary data and figures**

Figure S1. Zone of growth inhibition assays to test antibiotic sensitivity of B728a ∆*mexB* complementation variants. Mean values displayed as column height. For each compound tested, means marked with the same letter do not differ at p = 0.01 using Tukey’s HSD test. The lower limit of measurement is 6 mm (dotted line).

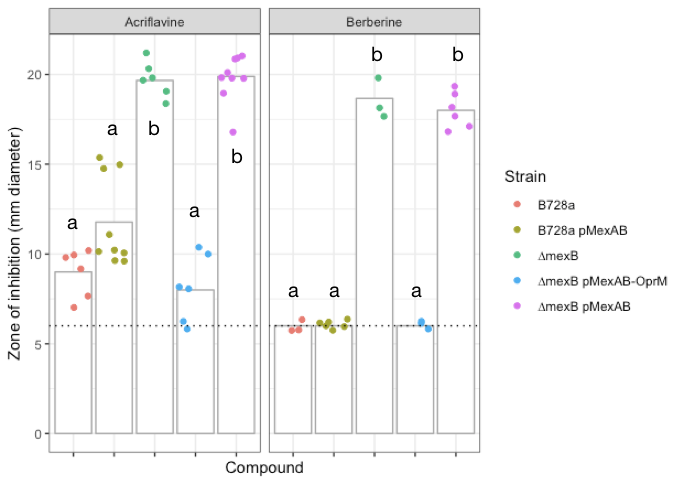

Figure S2. Fitness of transposon insertional mutants in the operon *Psyr_2282-3* in both a WT background and in cells in which *mexAB-oprM* has been disrupted. Fitness is calculated as the log_2_ change in relative insertion strain barcode abundance for a given gene.

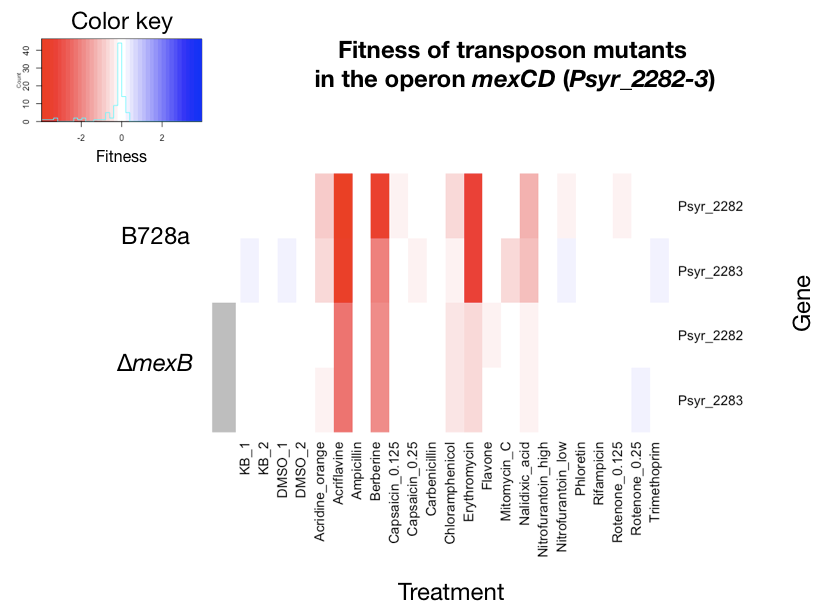

Figure S3. Fitness of transposon insertional mutants in the operon *Psyr_0917-8* in both a WT background and in cells in which *mexAB-oprM* has been disrupted. Fitness is calculated as the log_2_ change in relative insertion strain barcode abundance for a given gene.

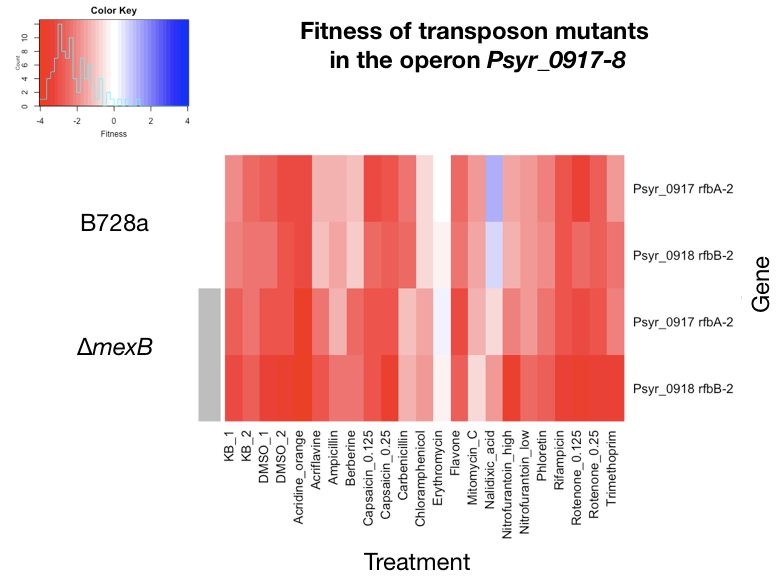

Figure S4. Fitness of transposon insertional mutants in the operon *Psyr_1754-9* in both a WT background and in cells in which *mexAB-oprM* has been disrupted. Fitness is calculated as the log_2_ change in relative insertion strain barcode abundance for a given gene.

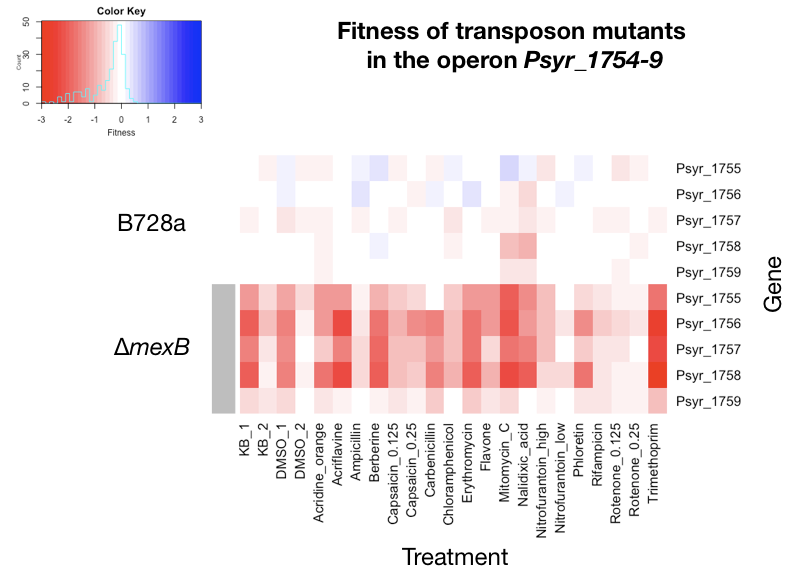
Figure S5. Structures of (A) acridine orange and (B) proflavine (top) / acriflavine (bottom). Acriflavine is typically sold as a mixture of 3,6-Diamino-10-methylacridinium chloride (acriflavine) with 3,6-diaminoacridine (proflavine) (www.sigmaaldrich.com).

A.

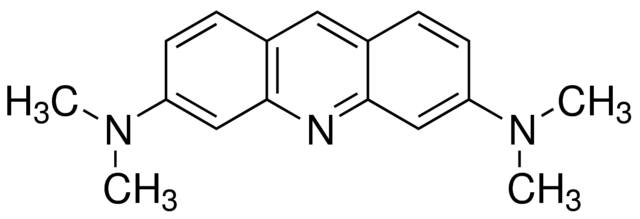

B.

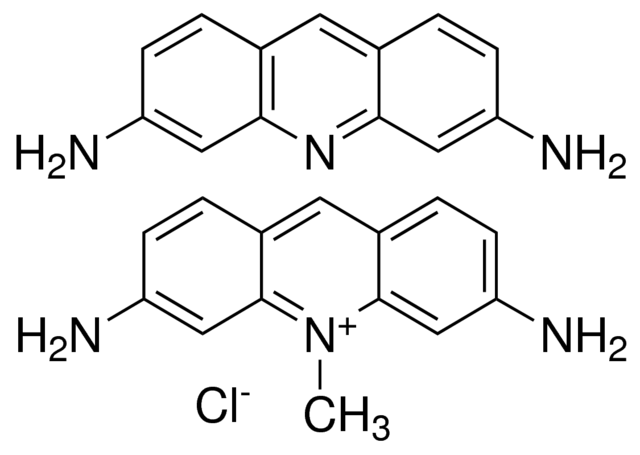

Table S1. Characteristics of the barcoded *mariner* transposon libraries in B728a WT and ∆*mexB* backgrounds. Genic strains contain insertions within the central 10 – 90% of the gene; only these insertions are used to calculate fitness. The total number of B728a genes = 5,216.

| **Library** | **B728a WT** | **B728a ∆*mexB*** |
| --- | --- | --- |
| Mapped barcodes | 281,419 | 237,285 |
| Genic strains | 169,826 | 143,498 |
| Median insertions / gene | 21 | 16 |
| Genes with enough reads | 4,296 | 4,255 |

Table S2. Antimicrobial compounds used in growth assays of the barcoded *mariner* transposon libraries. For compounds with known (and different) MIC values for the WT and ∆*mexB* genotypes, ¼ MIC values were used. Otherwise, the same concentration was used for both libraries. DMSO controls were used at concentrations representative of the highest solvent concentration used.

| Compound | Solvent | µg/ml for B728a | µg/ml for ∆*mexB* |
| --- | --- | --- | --- |
| DMSO_1 |  | 1% | 0.42% |
| DMSO_2 |  | 1.25% | 1.25% |
| Acridine orange | H_2_0 | 62.5 | 3.125 |
| Acriflavine | H_2_0 | 7.75 | 1.25 |
| Ampicillin | H_2_0 | 15.6 | 0.775 |
| Berberine | DMSO | 250 | 15.6 |
| Capsaicin | DMSO | 250 | 250 |
| Capsaicin | DMSO | 125 | 125 |
| Carbenicillin | H_2_0 | 250 | 3.1 |
| Chloramphenicol | EtOH | 12.5 | 1.55 |
| Erythromycin | EtOH | 25 | 6.25 |
| Flavone | DMSO | 125 | 125 |
| Fusaric acid | DMSO | 250 | 15.6 |
| Mitomycin C | H_2_0 | 0.625 | 0.05 |
| Nalidixic acid | H_2_0 | 12.5 | 0.775 |
| Nitrofurantoin | DMSO | 250 | 62.5 |
| Nitrofurantoin | DMSO | 125 | 31.25 |
| Phloretin | DMSO | 250 | 125 |
| Rifampicin | MeOH | 100 | 100 |
| Rotenone | DMSO | 250 | 250 |
| Rotenone | DMSO | 125 | 125 |
| Trimethoprim | H_2_0 | 125 | 15.6 |

Table S3. Shared amino acid identity between *P. syringae* B728a Psyr_2482-5 and *P. aeruginosa* PAO1 MuxABC-OpmB. Amino acid sequence homology was calculated using NCBI blastp. “Positives” includes amino acids having similar chemical properties.

| *P. syringae* B728a | *P. aeruginosa* PAO1 | Gene | A.A. identity | Positives |
| --- | --- | --- | --- | --- |
| Psyr_2482 | PA2528 | *muxA* | 219/420 (52%) | 279/420 (66%) |
| Psyr_2483 | PA2527 | *muxB* | 731/1022 (72%) | 875/1022 (85%) |
| Psyr_2484 | PA2526 | *muxC* | 611/1039 (59%) | 777/1039 (74%) |
| Psyr_2485 | PA2525 | *opmB* | 288/459 (63%) | 350/459 (76%) |

Strains and plasmids

| Strains | | |
| --- | --- | --- |
| Organism | Description | Source |
| *E. coli* TOP10 | For general cloning | Invitrogen |
| *E. coli* XL1-Blue | For general cloning | QB3 Macrolab |
| *E. coli* S17-1 | Conjugation donor strain | (54) |
| *E. coli* WM3064 | Strain APA752; barcoded *mariner* transposon vector (Kan^R^) in *E. coli* conjugation strain | (29) |
| *P. syringae* B728a | Wild type strain (Rif^R^) | (43) |
| *P. syringae* B728a | Whole genome barcoded *mariner* transposon library (Rif^R^ Kan^R^) | This work |
| *P. syringae* B728a | Whole genome barcoded *mariner* transposon library (Rif^R^ Kan^R^), in ∆*mexB* genotype | This work |
| *P. syringae* B728a | ∆*mexB* (Rif^R^ Kan^R^) | This work |
| *P. syringae* B728a | ∆*mexB* (Rif^R^) – cured of pFLP2Ω | This work |
| *P. syringae* B728a | p519-MexAB (Rif^R^ Kan^R^) | This work |
| *P. syringae* B728a | ∆*mexB* p519-MexAB (Rif^R^ Kan^R^) | This work |
| *P. syringae* B728a | ∆*mexB* p519-MexAB-OprM (Rif^R^ Kan^R^) | This work |
| *P. syringae* B728a | ∆*mexF* (Rif^R^ Kan^R^) | This work |
| *P. syringae* B728a | ∆*mexB* ∆*mexF* (Rif^R^ Kan^R^) | This work |
| *P. syringae* B728a | ∆*muxB* (Rif^R^ Kan^R^) | This work |
| *P. syringae* B728a | ∆*mexB* ∆*muxB* (Rif^R^ Kan^R^) | This work |
| *P. syringae* B728a | ∆*mexK* (Rif^R^ Kan^R^) | This work |
| *P. syringae* B728a | ∆*mexB* ∆*mexK* (Rif^R^ Kan^R^) | This work |

| Plasmids | | | |
| --- | --- | --- | --- |
| Plasmid name | Description | Antibiotic | Reference |
| pKD13 | Source of kanamycin resistance | Kan | (51) |
| pT*sacB* | Suicide plasmid to introduce DNA into *P. syringae* | Tet | (52) |
| pFLP2Ω | Contains Flp recombinase | Spec | (47) |
| p519*ngfp* | *lac* promoter and *pnpt2* in front of *gfp* | Kan | (46) |
| p519-MexAB | *mexAB* and upstream 1161 bp, replacing *gfp* in p519*ngfp* | Kan | This work |
| p519-MexAB-OprM | *mexAB-oprM* and upstream 1161 bp, replacing *gfp* in p519*ngfp* | Kan | This work |
| pT:2968-kan | To delete *mexF*, contains *Psyr_2968* flanking regions bordering kan^R^ cassette, inserted into SmaI site of pT*sacB* | Tet Kan | This work |
| pT:2483-kan | To delete *muxB*, contains *Psyr_2483* flanking regions bordering kan^R^ cassette, inserted into SmaI site of pT*sacB* | Tet Kan | This work |
| pT:0346-kan | To delete *mexK*, contains *Psyr_0346* flanking regions bordering kan^R^ cassette, inserted into SmaI site of pT*sacB* | Tet Kan | This work |

Primers.

Primers used to construct MexAB (*Psyr_4007-8*) and MexAB-OprM (*Psyr_4007-9*) expression plasmids. Restriction cut sites are lowercase.

| Name | Sequence |
| --- | --- |
| XbaI 4007 F | TAAtctagaTTTTCCTGCGGGGCAATACT |
| EcoRI 4008 R | ATgaattcGAACGCGGTAACTGCAAGTG |
| EcoRI 4009 R | ATgaattcCACTGATCGCAGGTGTGTTT |

Bold sequence complements FRT-Kan sequence for splicing by overlap extension protocol. The primers used for TnSeq mapping and BarSeq amplification are described (29).

| Name | Sequence |
| --- | --- |
| FRT-KanF | GTGTAGGCTGGAGCTGCTTC |
| FRT-KanR | ATTCCGGGGATCCGTCGACC |
| 4008 up F | AGCACAGCCCTATACCCTGA |
| FRT 4008 up R | **GAAGCAGCTCCAGCCTACAC**TTACTCCCCTTTGCTGCCTG |
| FRT 4008 down F | **GGTCGACGGATCCCCGGAAT**TGCAGTTACCGCGTTCATTC |
| 4008 down R | AGGAAGGTCAGGTTGCTGTC |
| 2968 up F | GGGATGAATTCACCGGTCGT |
| FRT 2968 up R | **GAAGCAGCTCCAGCCTACAC**CGGACGAGTCCCTTAGCCGC |
| FRT 2968 down F | **GGTCGACGGATCCCCGGAAT**GAGTGCGTACAAAGTCTTCATCCCG |
| 2968 down R | TTGTCGTAGTCGCTGAACGC |
| 2483 up F | GCAGCTATCAAGTGGCGTTG |
| FRT 2483 up R | **GAAGCAGCTCCAGCCTACAC**GGTTTTTCCACCTTGCCGAG |
| FRT 2483 down F | **GGTCGACGGATCCCCGGAAT**GTGAGCTTTCGCTTGTTGCC |
| 2483 down R | GAGCGGTCCATCGCGATATT |
| 0346 up F | CAAAGTGCGCGAAGTAACCC |
| FRT 0346 up R | **GAAGCAGCTCCAGCCTACAC**TGACTGGCGACGGTGAATAC |
| FRT 0346 down F | **GGTCGACGGATCCCCGGAAT**GGCTGCTGACGCGGTAATC |
| 0346 down R | CTGCGAAGTGGTGCAGCAAA |

Primers used to confirm deletion strains.

| Name | Sequence |
| --- | --- |
| 4008 check F | CAGTGGCTGTAGCAAGAAGGA |
| 4008 check R | TTTGTTGCGCACTGAACAGC |
| 2968 check F | GTACTGGAGCAGCCAGTCAA |
| 2968 check R | CCAGCAGGACCAGAAAGTCC |
| 2483 check F | TTCCAGGAAGGGCAGATGGT |
| 2483 check R | AGGTGCATGATCGCAAAGGT |
| 0346 check F | ACGTTGCTCGATAACCCGAA |
| 0346 check R | TCCGAACGTTTCGCAGAGAT |
